# The response to drought-stressed host plants varies among herbivorous mite populations from a climate gradient

**DOI:** 10.1101/2021.10.21.465244

**Authors:** Alain Migeon, Philippe Auger, Odile Fossati-Gaschignard, Ruth A. Hufbauer, Maëva Miranda, Ghais Zriki, Maria Navajas

## Abstract

Drought associated with climate change can stress plants, altering their interactions with phytophagous arthropods. Drought not only impacts cultivated plants but also their parasites, which in some cases are favored by drought. Herbivorous arthropods feeding on drought-stressed plants typically produce bigger offspring and develop faster. However, it is unclear how much responses to drought stress differ among populations of herbivore species. Here, we evaluate variability among populations of a major agricultural pest, the two spotted spider mite, *Tetranychus urticae*, in response to drought stress. We compare key life history parameters of twelve populations that originate from climates ranging from wet and cool Atlantic locations to medium to dry hot Mediterranean locations. We evaluated how plant drought stress affects four life history traits: development time, fecundity, sex-ratio and emigration rate in an experiment comparing well-watered and drought-stressed bean plants. Mites feeding on drought-stressed plants developed faster and attempted to leave leaves less often, and young females were more fecund. The mites from wet temperate climates exhibited greater plasticity between the two water regimes than mites originating from dryer and hot climates, suggesting that the climate in the area of origin influences mite response to drought.

## Introduction

“A Quarter of Humanity Faces Looming Water Crises”, warned The New York Times of August 6^th^, 2019 (Sengupta and Cai, 2019). Extreme climatic events have already increased in frequency due to climate change, and these increases are forecast to continue (IPCC, 2021). Low water availability not only impacts cultivated plants by enhancing crop yield vulnerability to climatic events (Olesen et al., 2011) but also plant pests, which can be favored by drought (see Hamann et al., 2021 for a review). Thus, to support agricultural production in the face of intensified drought, it is important to understand how pests respond to plant water availability.

Positive effects of drought stressed plants on herbivores are likely driven by changes in plant physiology (Chaves et al., 2003) including shifts in the amino-acids and free sugar balances in drought-stressed plants (Hummel et al., 2010; Showler, 2013). These shifts are thought to drive changes of life history traits. For example, drought stress in beans increased oviposition by the bug *Orius insidiosus* (Seagraves et al., 2011) and drought stress in the grass *Holcus lanatus* increased offspring and rates of emergence of the moth *Spodoptera littoralis*, increasing their fitness overall relative to well-watered plants (Walter et al., 2012).

Among plant pests, the responses of spider mites (Acari: Tetranychidae) to drought stress have been studied in field and laboratory settings, leading to diverse and sometimes divergent results. Drought stress of soybeans led to faster development and thus increased density of the spider mite *Tetranychus turkestani* (Nikolova et al., 2014). This pattern was also observed in *Tetranychus urticae* and *Oligonychus pratensis* on maize (Chandler et al., 1979), and in a mixed mite population of *Tetranychus pacificus* and *Panonychus citri* on almond (Youngman and Barnes, 1986; Youngman et al., 1988). Gillman et al. (1999) observed that the damage caused by *T. urticae* increased on drought stressed buddleia plants. Ximénez-Embún et al. (2016, 2017a, 2017b) observed a global increase of performance of three important tomato mite pests, *Tetranychus evansi, T. urticae* and *Aculops lycopersici*, reared on drought-stressed tomato plants, especially for tomato-adapted strains in the case of T. urticae. This faster development can lead to apparent trade-offs in fecundity as reported by Van Petegem et al. (2016). In addition, higher fecundity of young female can also lead to lower fecundity of older females (Youngman et al., 1988). A non-linear response with an increase of density and fecundity of *T. urticae* was reported at an intermediate level of drought, and a linear increase of development rate at a severe drought stress regime (English-Loeb, 1989). In contrast, the opposite pattern was reported by Oloumi-Sadeghi et al. (1988) who observed a decrease of *T. urticae* abundance on drought-stressed soybean and by Sadras et al. (1998) for *T. urticae* on cotton.

These divergent results in how mites respond to drought-stressed plants might be explained in part by intraspecific variation among mite populations from different locations. In general, the degree to which different populations of phytophagous arthropods differ in their responses to abiotic factors is poorly documented, but evidence suggests populations from different environments may often differ. For example, populations of a leaf beetle, *Diorhabda carinulata*, differ in the timing of diapause introduction across their North American range, enabling them to persist in environments with long cold winters south to deserts with milder winters (Bean et al., 2012). Among the tetranychid mites, common garden experiments reveal a latitudinal gradient of life history traits (decrease in fecundity, shortening development time, increase in male proportion and in dispersal) from Western European core distribution of *T. urticae* to the northernmost part of the distribution area (Van Petegem et al., 2016). These studies (Bean et al., 2012; Van Petegem et al., 2016) thus highlight the importance of intraspecific trait variation in response to climate.

Intraspecific variation of this nature can be evident as fixed differences among populations from different environments, often due to adaptation to the local environment or to differences in phenotypic plasticity in response to the environment. When environments are constant and predictable through time, fixed adaptive differences are more likely, whereas when environments vary strongly through time, phenotypic plasticity is more likely (reviewed in Olazcuaga et al., 2022). Thus, environments that are consistently arid may produce populations with less phenotypic plasticity than environments that can vary strongly in water availability throughout and across years.

Here, we evaluate how intraspecific genetic variation modifies responses to drought stress, focusing on populations of the two spotted spider mite, *T. urticae*, sampled from locations in Europe with distinct climates. From the review of literature, we hypothesized that on drought-stressed plants relative to well-watered plants, females from all populations (1) will develop faster; (2) will have higher fecundity; (3) will decrease their leaving rate; (4) will give rise to a more female biased progeny. We also hypothesized that populations from drier regions, which are likely to more consistently experience drought-stressed plants, may exhibit less plasticity in these traits than populations from wetter more temperate regions that may experience only occasional drought stress. To test these hypotheses, we evaluated how mites from 12 sampling locations that differ in climate responded to drought stress in their host plants. We examine the development time of females, their fecundity, the sex ratio of their progeny and the leaving rate on drought-stressed and well-watered host plants. Finally, we relate the observed variation in these four life history traits to the climate of origin, and explore the potential adaptive value of different phenotypes.

## Material and methods

To evaluate the effects of drought stress on mite life history traits, we conducted two experiments. Experiment I focused on development time of females on drought-stressed and well-watered plants, as shorter development time can increase population growth rate, and thus is an important life history trait linked to population performance. In experiment II, we evaluated how fecundity and sex ratio of offspring respond to drought stress, possibly through a temporal shift in their life span, and also measured attempts of mites to depart from experimental arenas.

### 1 Mites

#### Biological features

*Tetranychus urticae* is a cosmopolitan highly polyphagous pest recorded from 124 countries and 1169 plant species (Migeon and Dorkeld, 2019). It has two different color morphs (green and red), which can be observed in sympatry (Auger et al., 2013). These color morphs do not vary systematically in geographic distribution or physiological parameters. It is an arrhenotokous species, meaning that fertilized eggs from diploid females produce diploid females, while unfertilized eggs produce haploid males (Helle and Bolland, 1967). Sex ratio is usually female biased (between 60 to 70% of females), and shifts in sex ratio are one way populations respond to changing environments (Crozier, 1985).

*Tetranychus urticae* feeds by piercing leaf parenchyma cells with its stylet and sucking out the cell contents (Tomczyk and Kropczyńska, 1985). It develops from egg to adult in 10 days (Sabelis, 1981) at 25°C and is, as many Tetranychidae species, commonly reared between 20 and 25°C (Crooker, 1985). The embryonic (egg) development represents almost a half of the total duration of the development from egg laying to the adulthood. It is followed by three mobile stages interspersed by three immobile molting stages (Sabelis,1981). At 25°C, one female can lay a mean of 70 to 130 eggs throughout her reproductive life of approximately two weeks. Most eggs are laid in the first week with peak fecundity at three days after emergence (Sabelis, 1981).

#### Origin of mites

We sampled *T. urticae* from 12 locations in Europe that represent a wide range of climatic conditions (Migeon et al., 2019) where the mite can develop (Litskas et al., 2019) (Table 1, Figure 1). At each location, 30-50 females were collected to found laboratory populations as described below.

**Figure 1.**
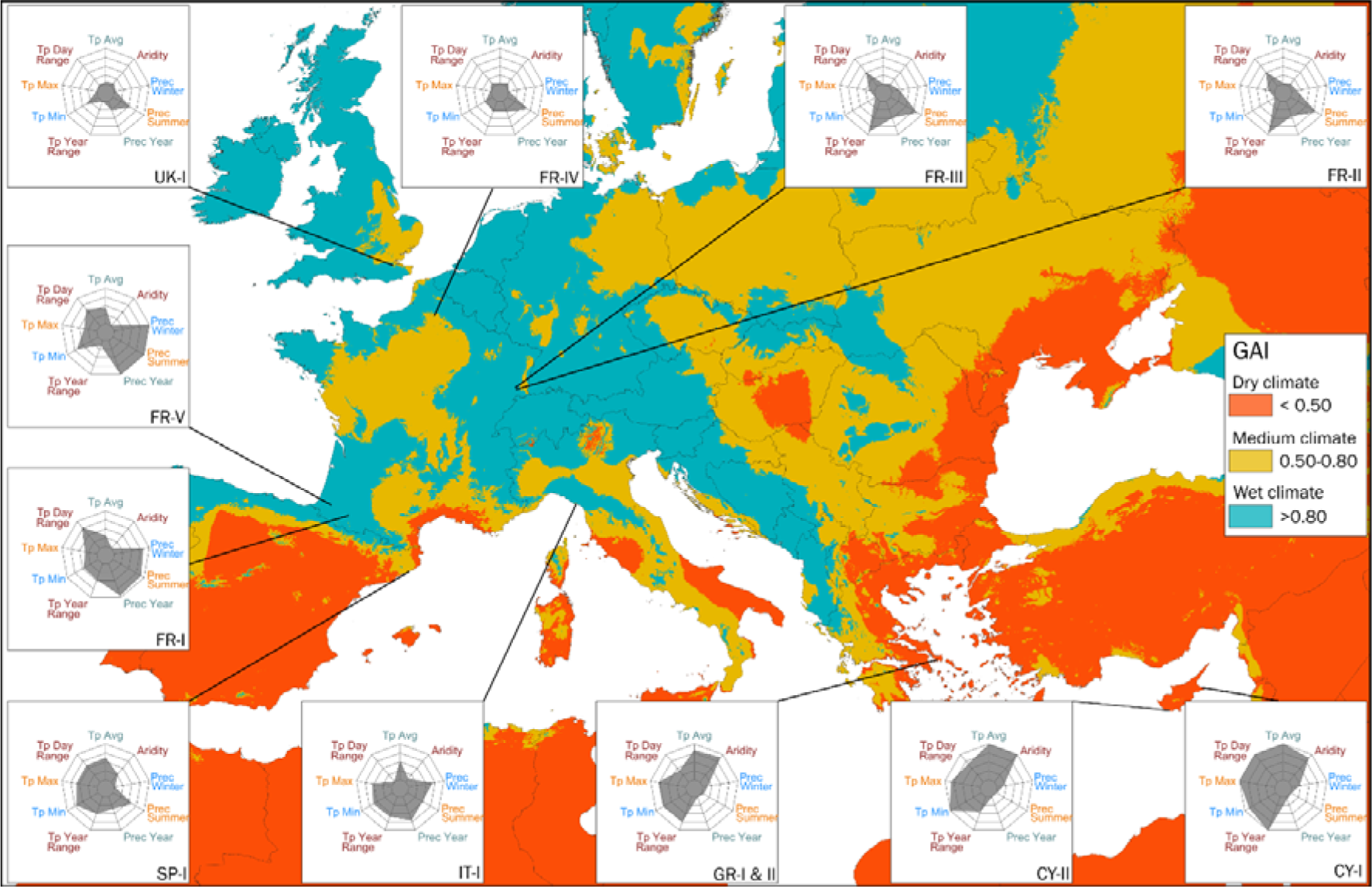
Map of the locations sampled. The background color is scaled on Global Aridity Index (GAI) (Trabucco and Zomer, 2019). The radar charts display the values of some important climatic variables: (from top, clockwise) annual temperature average (Tp Avg), 1/global aridity index (Aridity), precipitations of the coldest quarter (Prec Winter), precipitations of the warmest quarter (Prec Summer), total year precipitations (Prec Year), temperature annual range (Tp Year Range), minimal temperature of the coldest month (Tp Min), maximal temperature of the warmest month (Tp Max), mean diurnal range (Tp Day Range).

**Table 1.** Characteristics of the populations of *Tetranychus urticae* used in the experiments. DD: decimal degree.

| A. Location, climate and collected plant. |  |  |  |  |  |  |  |  |
| --- | --- | --- | --- | --- | --- | --- | --- | --- |
| Mite population | Mite body color | Global Aridity Index | Dryness | Country | Locality | Latitude (DD) | Longitude (DD) | Collection plant |
| CY-I | Red | 2433 | Dry | Cyprus | Paralimni | 35.050 | 33.990 | *Convolvulus arvensis* |
| CY-II | Green | 2176 | Dry | Cyprus | Kouklia | 34.693 | 32.578 | *Convolvulus arvensis* |
| FR-I | Red | 10725 | Wet | France | Livron | 43.227 | -0.139 | *Urtica dioica* |
| FR-II | Red | 8379 | Wet | France | Burnhaupt | 47.741 | 7.142 | *Urtica dioica* |
| FR-III | Green | 9003 | Wet | France | Guewenheim | 47.756 | 7.091 | *Urtica dioica* |
| FR-IV | Green | 7879 | Medium | France | Cappy | 49.928 | 2.783 | *Solanum tuberosum* |
| FR-V | Green | 11267 | Wet | France | Salies | 43.464 | -0.916 | *Urtica dioica* |
| GR-I | Green | 2387 | Dry | Greece | Karystos | 38.027 | 24.404 | *Malva sp* |
| GR-II | Green | 2387 | Dry | Greece | Karystos | 38.027 | 24.404 | *Xanthium italicum* |
| IT-I | Red | 7889 | Medium | Italy | Lerici | 44.086 | 9.889 | *Convolvulus arvensis* |
| SP-I | Red | 5289 | Medium | Spain | Palafolls | 41.671 | 2.758 | *Phaseolus vulgaris* |
| UK-I | Green | 7879 | Medium | United Kingdom | East-Mailing | 51.285 | 0.448 | *Urtica dioica* |

| B. Collection date and experimentation dates. |  |  |  |  |  |
| --- | --- | --- | --- | --- | --- |
| Mite population | Collection date | Experiment 1 | Days before exp. 1 | Experiment 2 | Days before exp. 2 |
| CY-I | 2016-06-02 | 2017-02-13 | 256 | 2017-04-06 | 308 |
| CY-II | 2016-06-01 | 2017-09-15 | 471 | 2017-10-27 | 513 |
| FR-I | 2017-07-11 | 2018-01-05 | 178 | 2018-05-07 | 300 |
| FR-II | 2017-09-02 | 2018-01-05 | 125 | 2018-05-07 | 247 |
| FR-III | 2017-09-02 | 2018-03-09 | 188 | 2018-06-01 | 272 |
| FR-IV | 2017-09-06 | 2018-03-09 | 184 | 2018-06-01 | 268 |
| FR-V | 2017-07-10 | 2018-03-30 | 263 | 2018-08-17 | 403 |
| GR-I | 2016-06-09 | 2017-09-15 | 463 | 2017-10-27 | 505 |
| GR-II | 2016-06-09 | 2017-10-20 | 498 | 2018-01-12 | 582 |
| IT-I | 2015-09-16 | 2017-01-18 | 490 | 2017-04-06 | 568 |
| SP-I | 2015-10-27 | 2017-01-18 | 449 | 2017-06-02 | 584 |
| UK-I | 2016-09-25 | 2017-10-20 | 390 | 2018-01-12 | 474 |

#### Mite rearing

Mite stock populations were maintained separately by collection site on detached bean leaves (*Phaseolus vulgaris* cv Contender), commonly used as a “neutral” plant (Sabelis, 1981; Fellous et al., 2014; Sousa et al., 2019). Leaves were placed on moist cotton blanket in double-bottom plastic boxes (13.5 x 9.5 x 5 cm) with water reservoir and maintained in growth chambers at 21 ± 1°C, 60 ± 10% RH with a photoperiod of L/D 16/8 h which allows the development of one generation on two weeks. Each population was maintained for at least six generations before being used in the experiments.

### 2 Climate data

Temperature, precipitation and bioclimatic variables for the 12 locations were retrieved from WorldClim (Fick and Hijmans, 2017) and the Global Aridity Index was taken from CGIAR (Trabucco and Zomer, 2019). We used monthly minimum, maximum and average temperatures to construct Gaussen climatograms (Supplementary Figure S1). The Bioclimatic variables gather a set of 19 synthetic variables describing the climate. The Global Aridity Index (GAI) was developed to quantify the precipitation availability over atmospheric demand (Trabucco and Zomer, 2019). This is a synthetic variable expressing the moisture availability for potential growth of reference vegetation. For the analysis we grouped a priori the 12 mite sampling locations in three clusters of dryness (dry, medium and wet) of four locations each (Table 1).

### 3 Plant material

#### Plant production

French bean (*Phaseolus vulgaris* cv. Contender) plants were grown from seeds in 2 L pots (diameter: 15 cm, height: 17 cm) (CEP, HR 17YPP) filled with 815 g (R.H. about 30%, measured with a soil moisture sensor HH150 (Delta-T Devices Ltd, Cambridge, UK)) of peat mix (Huminsubstrat N2, Neuhaus, Klasmann-Deilmann, Geeste, Germany). For each experiment, two sets of pots were differentiated according to the watering regime: well-watered plants (high water regime) were watered to saturation every day and drought-stressed plants (low water regime) were watered only once at sowing time with 200 ml of water. Pots were first kept in a regulated greenhouse with additional light when necessary (25 ± 7°C / 40 ± 30% RH) for ten days after sowing.

#### Drought stress maintenance and assessment

Ten days after sowing, bean seedlings that had two expanded cotyledons were transferred to a climate chamber where they were watered differentially according to water treatment. Light was provided by agro red and blue LED lamps (Philipps Green Power LED). We manipulated water availability differently in our two experiments. Well-watered plants were maintained with a soil moisture above 45% (substrate saturation) and drought-stressed plants with a soil moisture between 10-8% (8% is over the wilting point). In experiment I, we used an automatically regulated drip irrigation system. Soil water content (RH) was measured and recorded using 5 moisture sensors (SM150 with GP2 Data Logger, Delta-T Devices Ltd, Cambridge, UK) in each watering treatment and linked to DeltaLINK 3.1.1 PC software (Delta-T Devices Ltd, Cambridge, UK) for setting up and downloading data from a GP2 station. In the well-watered treatment, when the average soil moisture dropped to 45%, each plant was automatically watered for 30 seconds (delivering 17 ml of water) by a drip. In the drought-stressed treatment, watering was activated (same duration and amount of water per watering event) when soil moisture dropped to 8% (see Supplementary Figure S2 as an example). In experiment II, plants were watered manually. In the well-watered regime, beans received 100 ml of water daily. In the drought-stressed regime, they received once 20 ml of water a single time when transferred to the climate chamber (10 days after sowing). Seven days later, we started watering these drought-stressed plants with 20 ml of water daily.

Water stress level of plants was assessed several times during the experiments. In both experiments, we measured leaf stomatal conductance a commonly used drought indicator (Verslues et al., 2006) using a leaf porometer (SC-1 Leaf Porometer, Decagon Devices, Inc., Pullman, WA, USA). For each population 10 plants were used to estimate stomatal conductance, 7 times between ages 11 days and 27 days after sowing. For young plants (less than 20 days after sowing) the sensor was placed head in the upper third of the seed leaf (the part closest to the petiole of the leaf), on the side of the leaf. For older plants (21-27 days after sowing) measurements were taken in the first trifoliate leaf, placing the sensor head on the side of the central leaflet (see Supplementary Figure S3). In experiment II, we also assessed, with the same frequency, the soil moisture using a soil moisture sensor (HH150, Delta-T Devices Ltd, Cambridge, UK) and by weighing the pot (seedling, pit mix and pot) using a balance (KSR1 Proline, Darty Ptc, UK) on 10 plants (the same ones along the experiment) per water regime (see Supplementary Figure S4).

### 4 Experimental design

Due to restricted space available, each population was assayed separately in random order. This experimental constraint means that comparisons among populations can only be done with caution. We address this below further where we describe the data analysis. All the experiments were conducted in a climate room with diurnal (25 ± 1°C) and nocturnal (23 ± 1°C) temperatures and using a light cycle of L/D 16/8 h. Relative humidity was maintained at 50 ± 20% RH using a dehumidifier (Rexair 2500T, Rexair, 95330 Domont, France).

#### Experiment I (Figure 2A)

**Figure 2.**
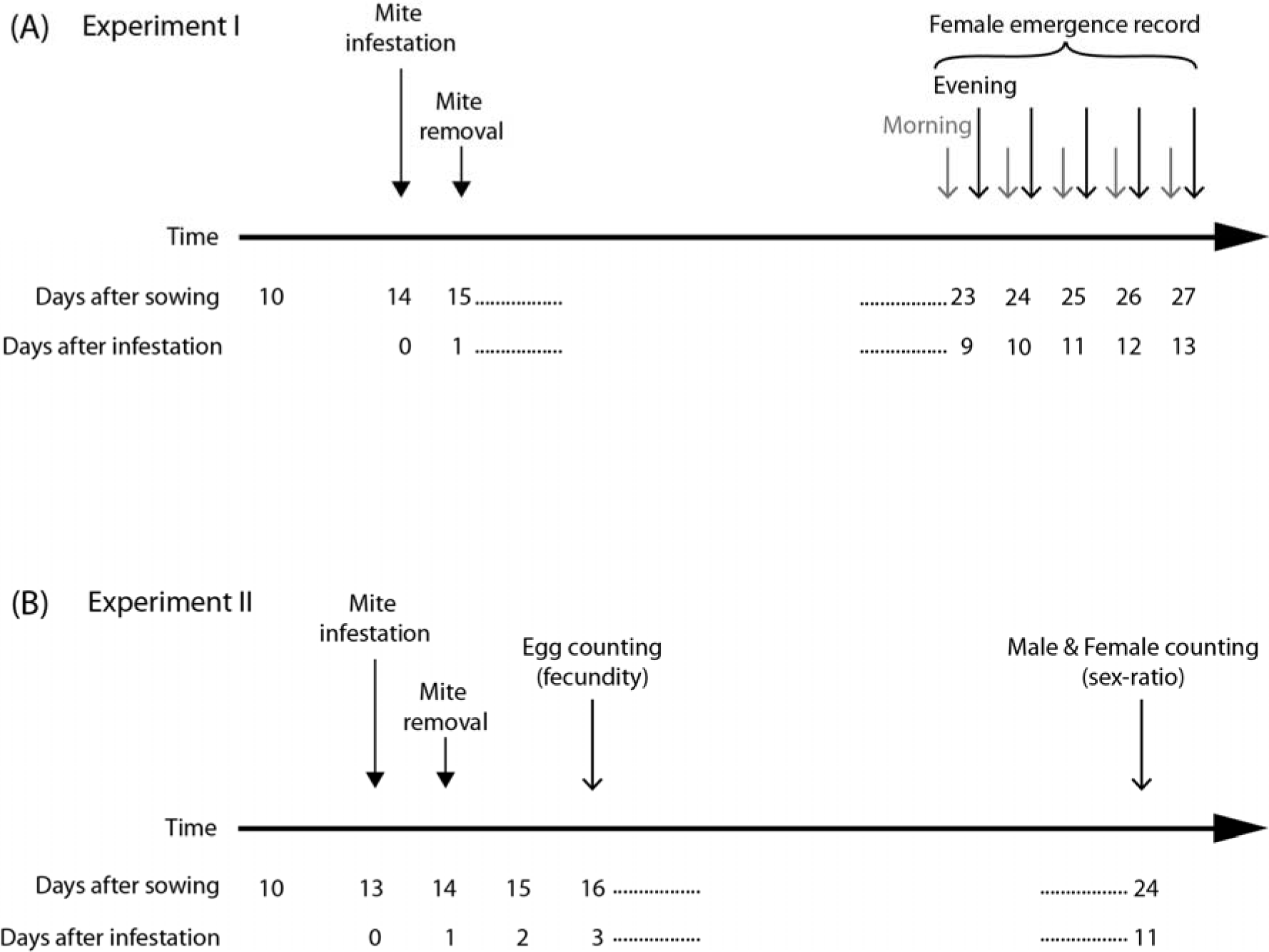
Timing of assessment of development time in female mites in Experiment I (A); timing of assessment of fecundity and sex ratio in Experiment II (B).

Experiment I was designed to estimate the development time of mites. Well-watered and drought-stressed plants with two expanded cotyledons were randomly arranged in the climate chamber and then infested with mites 14 days after sowing, at point steady stress conditions were reached in plants assigned to the drought-stressed treatment (Supplementary Figure S3).

Plants were infested by gently transferring 10 females of unknown age with a fine camel hairbrush, five per each cotyledon, into a squared arena built on leaves upper surface (Supplementary Figure S6). Females placed in these arenas were allowed to lay eggs for 24 hours and then removed (Figure 2A) using a Rena vacuum. Twelve to 15 replicates (12-15 plants) per water regime were performed for each mite population. From day 9 to 13 after infestation, newly emerged adult females were recorded and then removed twice a day, at 8 am and 4 pm (Figure 2A), using a stereomicroscope Leica EZ4 (Leica microsystems CMS GmbH, Wetzlar, Germany).

#### Experiment II (Figure 2B)

Experiment II was designed to assess the effect of plant drought stress on female fecundity, leaving rate and sex ratio of the progeny. Plants were produced with the same protocol than experiment I and transferred in the climate chamber 13 days after sowing. Then, they were infested with T. urticae females of known age. The mites used were transferred from plants under the same watering regime and monitoring as the treatment for which they were used (see Supplementary Figure S5).

A first batch of plants was infested with three-day-old females and six days later, a second batch (plants of the same age – 13 days after sowing) with nine-day-old females. The procedures for plant infestation and mite confinement were the same as those mentioned in experiment I. Females were allowed to lay eggs for 24 h and subsequently removed using a Rena vacuum. At this time, individuals were recorded as living, dead or drowned in the oily barrier. The mites found in the barrier were recorded to estimate attempts to leave the patch of leaf. Female fecundity was assessed by counting eggs two days after females were removed from arena with the aid of a stereomicroscope Leica EZ4 (Leica Microsystems, Weltzar, Germany). The eggs were kept on plants until hatching, and offspring allowed to develop to adulthood. The sex-ratio of newly emerged adult mites was then assessed 11 days after mite infestation.

### 5 Data analysis

All data analyses were conducted using R 3.5 (R Core Team, 2018). Graphics were produced using the library ggplot2 (Wickham, 2016).

The development time of mites was calculated by using a logit regression (library MASS, Venables and Ripley, 2002) to determine the time of 50% of emergence of adults. For each population, an ANOVA (Chi^2^ model) was used to test the effect of watering regime on development time.

The effect of watering regime on the other life history traits (fecundity, leaving rate and sex-ratio) was investigated with one t-test for each trait per population. This method was preferred to a common analysis because all the populations were assayed separately. Because escape rate (expressed as number of escaped females per plant / number of females deposited per plant) and sex-ratio (expressed as number of female per plant / (number of females per plant + number of males per plant)) are both limited from 0 to 1, they were respectively transformed to [arcsin(√(1-x))] and [arcsin(√x)] before analysis. Raw values are used in graphs and tables.

We conducted ANOVAs to examine trends of the change between watering regimes and dryness of the climate of origin. For each life history trait, we ran a model that included watering regime (two levels: well-watered or drought stress) and categorical dryness of the location of origin (three levels: wet, medium or dry), as determined from Global Aridity Index values (see Table 1) and the interaction between the two. The model was the following:

trait value ∼ watering regime x dryness

Population was not in the model because the populations in this case are the replicates, with only a single estimate for each watering regime. Normality of the residuals was tested by a Shapiro-Wilks test and homoscedasticity by a Levene test. Response variables for these ANOVAs included development time, fecundity of three-day-old and nine-day-old females, escape rate of three-day-old and nine-day-old females and sex ratio of the progeny of three-day-old and nine-day-old females.

To examine the degree of phenotypic plasticity in the measured traits, we calculated the differences observed in life history traits between watering regimes. A small difference indicates little plasticity, while a large difference indicates substantial plasticity. We used linear regressions to explore how the climate of origin affected the magnitude of the differences of the response in the life history traits between watering regimes. Specifically, we evaluated the relationship between the reduction of development time, the increase in fecundity, the reduction in leaving rate, and the change in the progeny sex-ratio with drought stress and the 20 climatic variables (19 Bioclimatic variables + Global Aridity Index) of each collection location.

## Results

### 1 Development time

Mites from all populations developed significantly faster when reared on drought-stressed plants (Figure 3 and 4A, Supplementary Table S1). The reduction in development time from the egg to adult ranged from 0.54 day (GR-II) to 1.35 day (FR-II).

**Figure 3.**
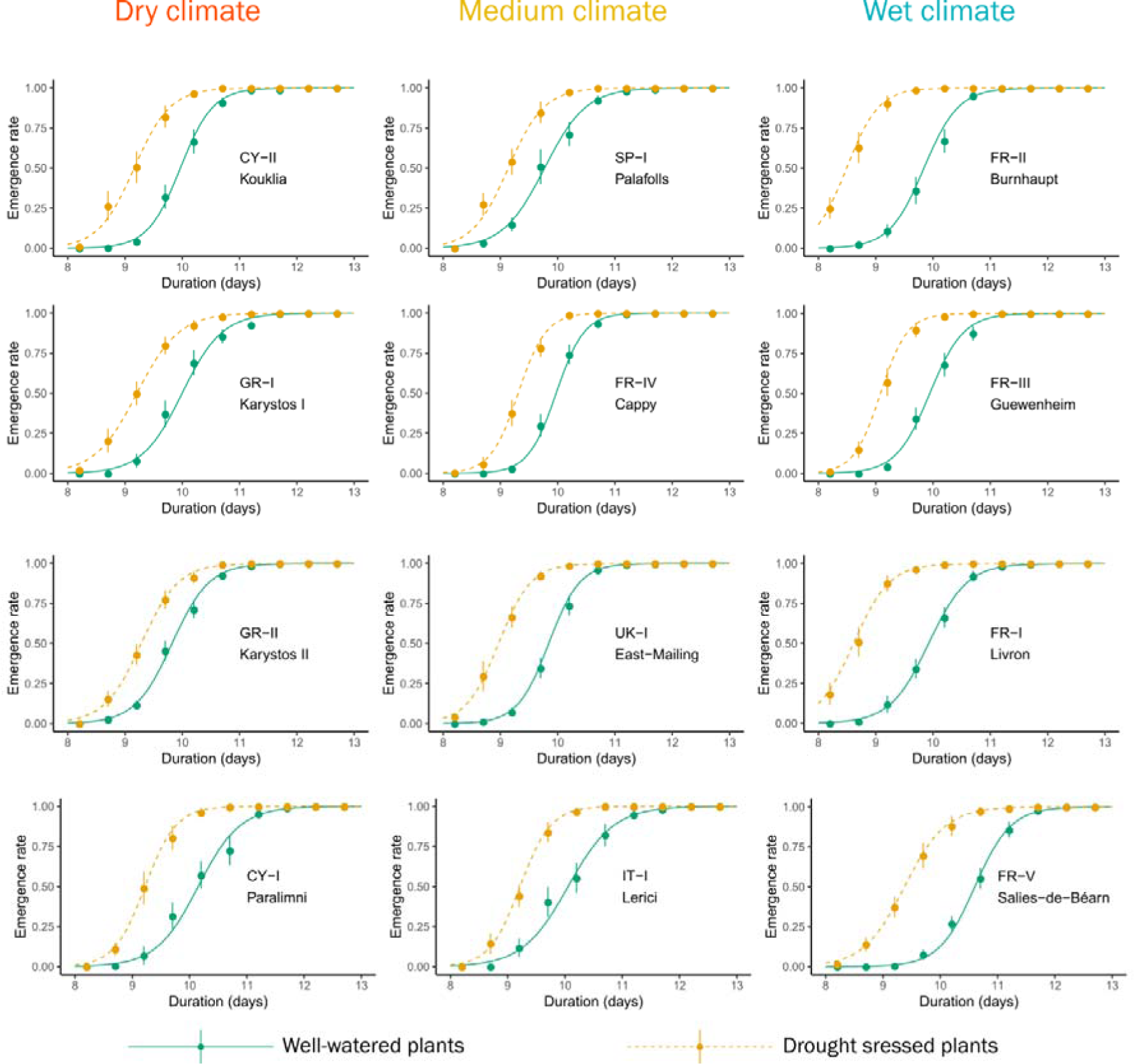
Development time from egg to adult of the twelve populations of *Tetranychus urticae* reared on well-watered and drought-stressed plants. Values are mean emergence rate of adults ± standard error at each observation point. Curves represent the logit regression.

**Figure 4.**
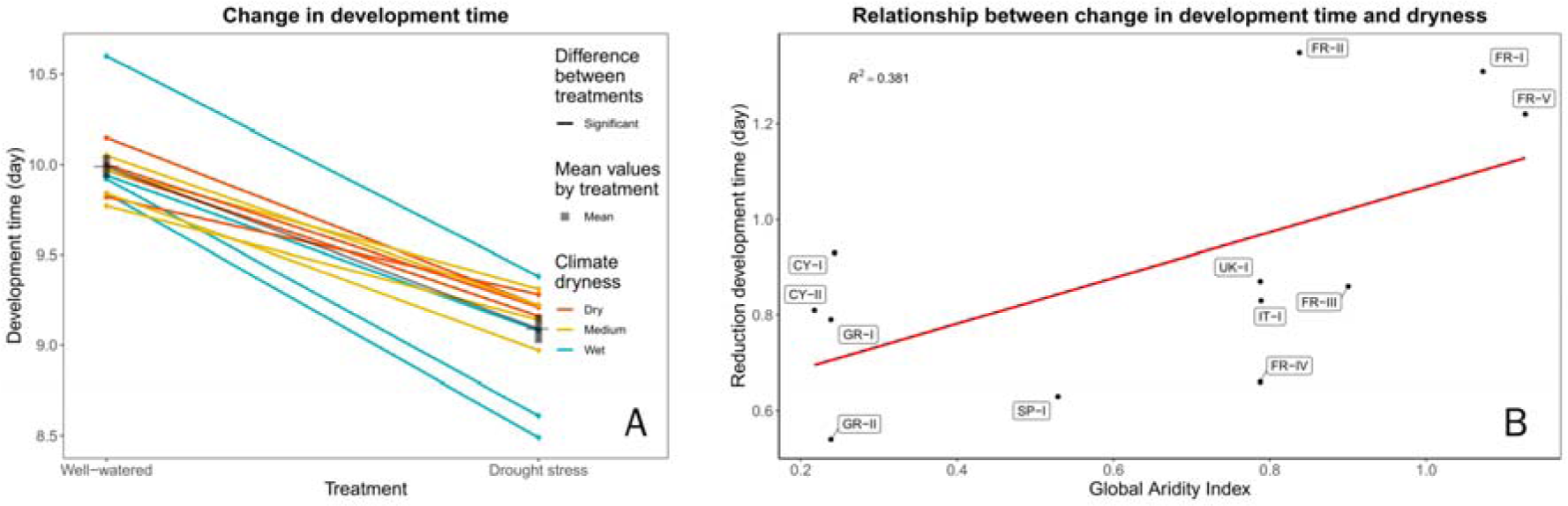
Change in development time of 12 populations of *Tetranychus urticae* between the two watering regimes. A: Development time on well-watered and drought-stressed plants. Values are mean development time (day) ± standard error for each population. B: Relationship between the change in development time and the Global Aridity Index (lower values indicate a drier climate) for each population, showing that mites from drier climates have smaller shifts in development time than mites from wetter climates.

Matching the individual population analyses, the ANOVA also showed a global significant effect of watering regime on development time (*F*_(1, 18)_ = 82.5, *p* < 0.001). The Global Aridity Index was a significant predictor of the reduction in development time (see Figure 4B and Table 2). Mites from locations with high summer humidity (high Global Aridity Index) responded more strongly to drought stress than mites from locations with low summer humidity. In addition, significant relationships between the reduction in development time and five others climatic variables were also observed (see Table 2). All but one of the climatic variables (BIO19, Winter Precipitations) refer to summer hydric local conditions that may be responsible for plant water stress, which is in line with the relationship obtained with the Global Aridity Index.

**Table 2.**
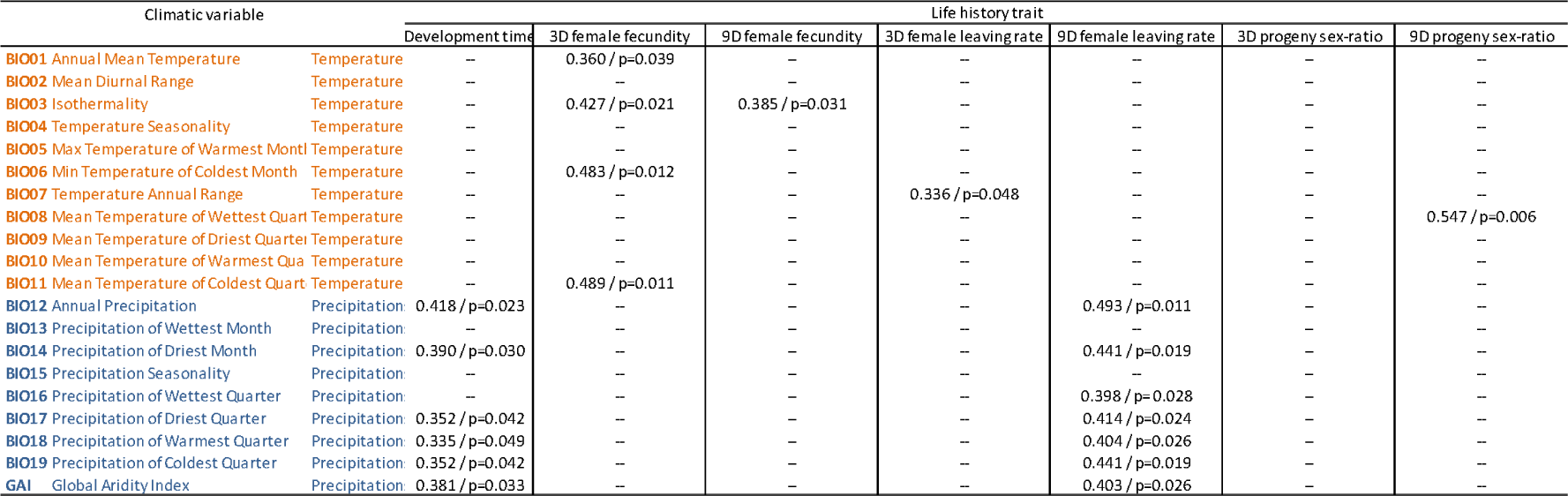
Regression coefficients between change in the life-history trait response of 12 populations of *Tetranychus urticae* between the two watering treatments and bioclimatic variables. R and p-values are reported when a significant relationship was observed.

### 2 Fecundity

The fecundity of three-day-old females increased on drought-stressed plants for 11 out of the 12 mite populations studied (Figure 5 and Supplementary Table S2). This increase was significant for seven populations: three from wet (FR-I, FR-II, FR-V), two from medium (IT-I, UK-I) and two from dry localities (GR-I, GR-II), and ranged from 0.26 (SP-I) to 2.62 (FR-II) eggs per female per day for a mean of 10.54 on well-watered plants and 11.52 eggs per female per day on drought-stressed plants. One population (CY-II) showed a significant decrease of fecundity on drought-stressed plants. The annual mean temperature was a significant predictor of the increase of fecundity on drought stressed plants (see Figure 6A and Table 2). In addition, significant relationship between the increase of fecundity and three other climatic variables, all linked to temperature and especially winter temperatures, were also observed (see Table 3). We also observed a correlation, on drought-stressed plants, between fecundity of three-day-old females and development time (r^2^ = 0.344, *p* = 0.045).

**Figure 5.**
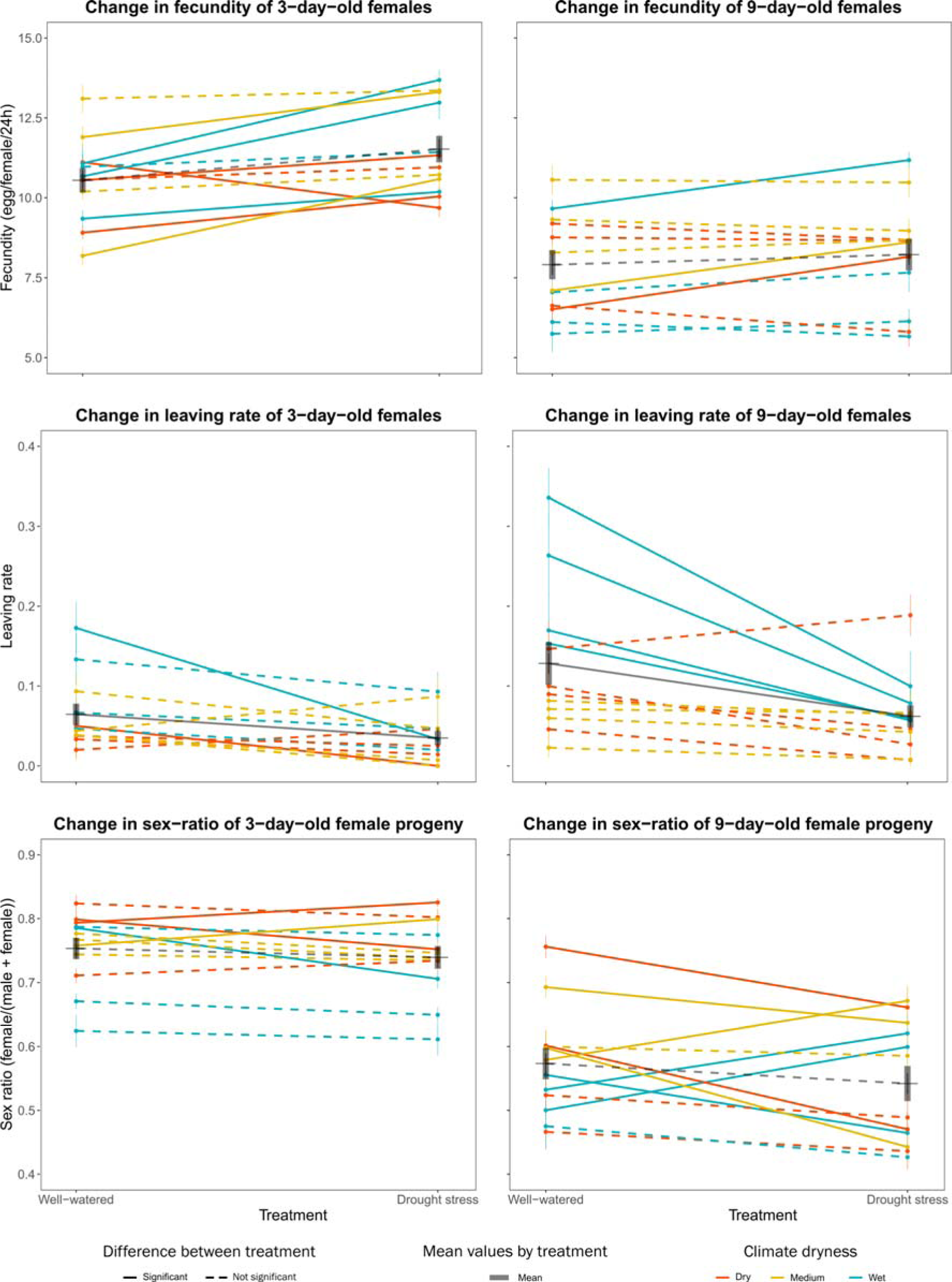
Variation in life history traits among the 12 populations of *Tetranychus urticae* studied, in response to plant watering regime. For each of the traits, values are mean ± standard error for each population.

**Figure 6.**
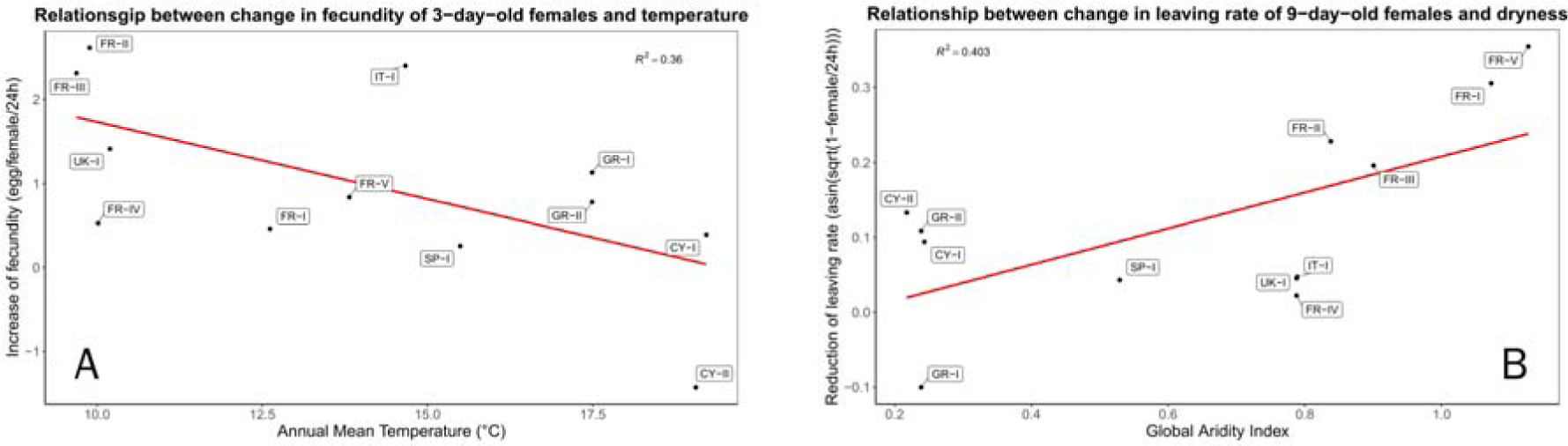
Relationships between changes in life history traits values between mites reared on well-watered or drought-stressed plants of 12 populations of *Tetranychus urticae*. A: Relationship between the change in the fecundity of 3-day old females and Annual Mean Temperature for each population. B: Relationship between the change in the leaving rate of 9-day old females and Global Aridity Index (lower values indicate a drier climate) for each population.

The fecundity of nine-day-old females increased on drought-stressed plants for six out of the 12 mite populations studied (Figure 5 and Supplementary Table S2). This increase was significant for three populations: one wet (FR-III), one medium (IT-I) and one dry location (GR-II). Only isothermality (BIO03) was a predictor of the increase of fecundity of the old females on drought-stressed plants (see Table 2). We also observed a correlation between fecundity of three-day-old and nine-day-old females reared on well-watered plants (r^2^ = 0.395, *p* = 0.029).

### 3 Leaving rate

Ten of the twelve populations studied showed a decrease in the leaving rate of three-day-old females on drought-stressed plants (Figure 5 and Supplementary Table S3) ranging from 1.2% (GR-II) to 13.9% (FR-II) females leaving per 24 h. However, results were significant for only two of them: one from a wet location (FR-II) and one from a dry location (CY-I). The average mite leaving rate from drought-stressed plants (3.5%) was approximately half that from well-watered ones (6.4%). The ANOVA showed a global effect of watering regime on leaving rate (*F*_(1, 18)_ = 5.0, *p* = 0.038). The temperature annual range (BIO07) was a predictor of the decrease of leaving rate of three-day-old females on drought-stressed plant (see Table 2).

The leaving rate of nine-day-old females was generally higher for mites exposed to well-watered plants. Eleven of the 12 populations studied showed a decrease in the leaving rate on drought-stressed plants (Figure 5 and Supplementary Table S4) ranging from 0.05% (FR-IV) to 23.6% (FR-V) and the four from wet localities (FR-I, FR-II, FR-III, FR-V) were significant. Mites attempted to leave drought-stressed plants (6.2%) half as often as they did from well-watered plants (12.9%). The ANOVA showed a global effect of watering regime (*F*_(1, 18)_ = 8.5, *p* = 0.009) and of dryness of the localities (*F*_(2, 18)_ = 6.9, *p* = 0.006) on leaving rate. The Global Aridity Index was a predictor of the decrease of leaving rate of nine-day-old females on drought stressed plant (see Table 2 and Figure 6B). Mites from locations with high summer humidity (high Global Aridity Index) responded more strongly to drought stress than mites from locations with low summer humidity. Each of the six other climate variables BIO12, BIO14, BIO16, BIO17, BIO18 and BIO19 showing significant relationship with the reduction of leaving rate on drought stressed plants represent precipitations variables and six of them indicate water availability in summer.

As observed for development time, mites originating from the four most humid locations (FR-I, FR-II, FR-III, and FR-V) showed higher differences in the leaving rate between the two water regimes and in line with this, the climatic variables related to precipitation and aridity were linked to the correlations with differences in the two water regimes.

### 4 Progeny sex-ratio

The sex ratio of the progeny of three-day-old females showed significant differences between water regimes in four populations (Figure 5 and Supplementary Table S4) but two of them represented an increase of female proportion: one dry (CY-I) and one medium location (SP-I) and the two other a decrease: one dry (CY-II) and one wet location (FR-V). The ANOVA showed significant differences (*F*_(2, 18)_ = 4.6, *p* = 0.024) related to climate aridity, but no related to water regime and the post hoc Tukey HSD test showed a significant difference between the sex-ratio of dry locations and wet locations with a higher proportion of females in dry locations regardless of the watering regime. No relationships were observed between climatic variables and change of sex-ratio of the progeny of three-day-old females.

The sex-ratio of the progeny of nine-day-old females showed significant differences between water regimes in eight populations but three of them represented an increase of female proportion: two from wet (FR-II, FR-III) and one from medium location (SP-I) and five a decrease: one from wet (FR-V), one from medium (UK-I) and two from dry localities (CY-I, GR-II) (Figure 5 and Supplementary Table S4). Only the Mean Temperature of the Wettest Quarter (BIO08) was a predictor of the change in sex-ratio. We also observed a general decrease of female proportion in the progeny of the oldest females. This phenomenon is the result of sperm availability in oldest females which decrease along time (Krainacker and Carey, 1990).

## Discussion

We have assessed 12 populations of mites originating from contrasted climatic condition, especially summer aridity index, combined with two water regimes. Mites developed faster, were generally more fecund and dispersed less when reared on drought stressed plants. These three factors could together allow mite populations on drought-stressed plants to grow more rapidly. Not all the mite populations tested responded equally, and differences between them depended on the climate conditions experienced in their area of origin.

### 1 Drought-stressed versus non-stressed plants

The shortening of development time for all the studied populations is in line with previous experiments conducted on spider mites (Chandler et al., 1979; Youngman and Barnes, 1986; Youngman et al., 1988; English-Loeb, 1989; Nikolova et al., 2014). Mites collected from different origins have similar development time when reared on well-watered plants, suggesting that mite development time on physiologically balanced plants depends only on the host-plants species and the temperature. Temperature is a well-known factor governing development time of ectotherms (Logan et al., 1976). However, Van Petegem et al. (2016) found a relationship between development time and latitude (which in their study corresponds to a thermal gradient). Variation in development time was driven by faster development in the northern edge of the mite’s distribution, where new populations settle. They interpreted their results as a combination of local adaptation and spatial selection. In our study, on well-watered plants we did not observe a relationship between development time and any of the climatic variables of the locations where the mite populations were collected. This might be because we deliberately collected our spider mite populations from locations with different climatic profiles that were still within the core climate conditions of the species (Litskas et al., 2019).

The increase of fecundity, along with shorter development time, confirms the previous reports of increased performance of T. urticae mites reared on drought-stressed plants (Chandler et al., 1979; Youngman and Barnes, 1986; Youngman et al., 1988; Ximénez-Embún et al., 2017a; Santamaria et al., 2018). Thus, the physiological changes that occur in drought-stressed plants are reliable changes. In tomato, these shifts in mite life history are likely linked to increased concentration of essential amino acids and free sugars (Ximénez-Embún et al., 2017a), which improved the nutritional value of drought-stressed plants. Similar changes may be occurring in drought-stressed bean plants.

On drought-stressed plants, the decrease of development time and the increase of fecundity of young females are concomitant, which reinforces the hypothesis of a general beneficial effect on the overall performance of mites on drought stressed plants. The higher fecundity of young females on drought stressed plants does not appear to come at a cost in fecundity of nine-day-old females. These results are not in accordance with Youngman et al. (1988) about *T. pacificus* on almond trees. They observed a shift in peak fecundity, in which an increase in fecundity in the first ten days on drought-stressed plant was counterbalanced by a decrease after ten days. Our experimental design did not allow us to observe females older than nine days and so does not rule out a later shift in fecundity. The correlation we observed between fecundity of three-day-old and nine-day-old females reared on well-watered plants suggests that our populations differ inherently in their reproductive output. These results are in line with Van Petegem et al. (2016) who reported variation in lifetime fecundity of mites for populations originated from the mite’s core distribution varying in a scale from 20 to 110 eggs per female and not linked to latitude or temperature. In our experiments, the combination of fecundity and sex ratio show an increase of 0.58 females per female per day for the three-day old female progeny on drought stressed plants (from 7.94 to 8.51) and a decrease of 0.08 females per female per day (from 4.53 to 4.45). This phenomenon could be interpreted as a trade-off, but it could also be, more likely, the result of a decrease of sperm availability as a result of the effectiveness of the only first mating in spider mites (Helle, 1967), in nine-day-old females on drought-stressed plants as a result of the higher production of females in three-day-old female progeny (Krainacker and Carey, 1990). Fecundity, dispersal and sex-ratio are often linked by complex relationships (Van Petegem et al., 2016) and the quality of the environment which in turn can shape the nutritional quality and the attractiveness of the host plants to the arthropods, also impacts these relationships. The higher leaving rate of the older females on well-watered plants may be linked to the lower performance of those same females on those plants with respect to fecundity and development time. However, in nature, the surrounding environment, mostly driven by the climate, may play a role by increasing or decreasing the risk of leaving a plant for a potential new plant.

### 2 The importance of climatic condition of sampled locations

Our study, by comparing populations from distinct climatic origins, reveals that changes in life history traits of mites feeding on drought-stressed plants depended on the climate of their area of origin (Figure 7). Though it was not feasible to run all the trials at the same time, the results show enough consistency to be interpreted together. Since all the females coming from the different locations were reared under identical conditions, it is reasonable to accept that observed variations resulted from genetic differentiation in the tested populations. Our main finding is that mites originating from wet to cool locations displayed a stronger response to drought stress. Their three life history traits linked to climate (development time, fecundity of young females and leaving rate of old females) changed more between drought-stressed and well-watered plants than did mites from the drier locations. The changes in phenotype with plant treatment of the mites from drier hotter locations were weaker, except for the change in fecundity of CY-II (the unique population with a decrease on drought-stressed plant) and the leaving rate of GR-I (the unique population with an increase on drought-stressed plants). These results support the hypothesis that populations from drier locations which are likely to more consistently experience drought-stressed plants, exhibit less plasticity in the traits involved in the response to drought stress than the populations from the wetter regions.

**Figure 7.**
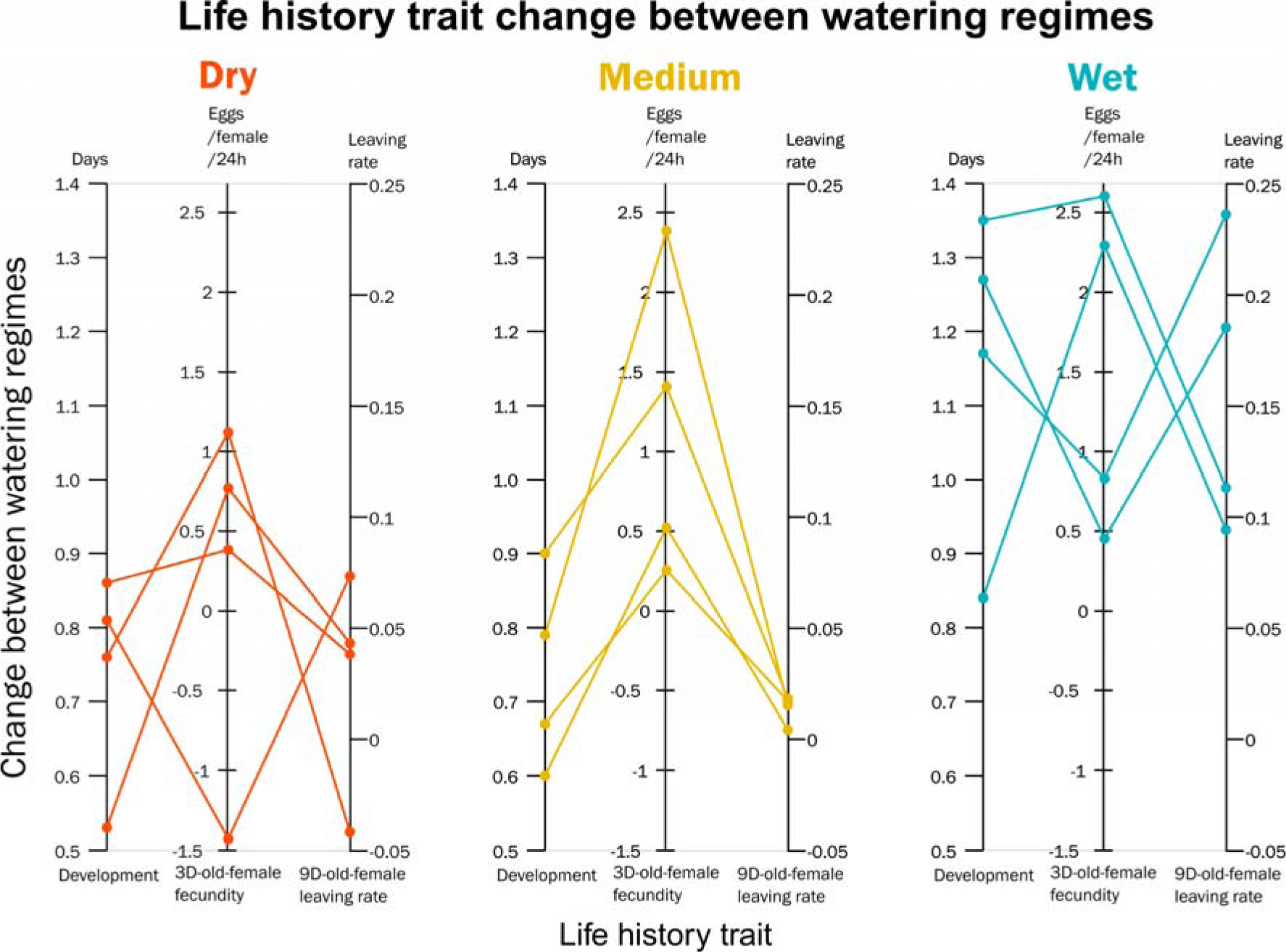
Differences between the two watering regimes for the three life history traits showing responses linked to climate (see Table 2): development time (DEV), fecundity of 3-day-old females (FEC3) and leaving rate of 9-day-old-females (LR9). The 12 populations of *Tetranychus urticae* studied are categorized by aridity of the location.

Previous study (Chen et al., 2020) highlighted that genetic variation for two closely related species *Tetranychus truncatus* and *Tetranychus pueraricola*, were associated with climatic parameters, mainly temperature and precipitation across China. For both species, genotype association was stronger with precipitation parameters together with the neuropeptide receptor NPR-9 gene adjacent genomic region. The NPR-9 affects foraging behavior and nutrient storage (Bendena et al., 2008) and as a consequence development time and fecundity. These studies (Chen et al., 2020 and Bendena et al., 2008) tend then to support that local adaptation to diverse levels of aridity could shape mite responses allowing them to adjust feeding behavior in accordance with native local climatic conditions and nutritional quality of the host plants.

### 3 Conclusion

Lehmann et al. (2020) underline the necessity of explore intraspecific variability of pests to forecast their response to climate warming. They highlighted four major categories of responses to climate warming: range changes, life history traits, population dynamics and trophic interactions. Our results, focused on the effects of drought, complement their analysis, and suggest that the changes observed in life history traits may act together with direct responses to temperature to enhance the rate of population growth, modifying the population dynamics of pests and potentially also the trophic interactions with their predators (see Litskas et al., 2019). The largest changes observed for populations from the wetter localities also reinforce the hypothesis of a more acute increase of crop losses in temperate areas (Deutsch et al., 2018).

However, our results also mitigate this general analysis. Indeed, populations originating from wet to cool locations (Alsace and Pays Basque) have not (or rarely) been submitted to drought-stressed host plants before, while populations from Cyprus and Greece had to face harsh climate and dryness half of the year. Our results, although not proving it, do suggest that the “over-reaction” of the French and British populations could have a cost, which might limit the response to drought of populations from dry locations. However, populations from drier climates may also have adapted to those conditions, and, in the absence of costs, their life history traits do not differ as much from those in control conditions in contrast with non-adapted populations. As stated by Hoffmann (2017), a genomic analysis would be necessary to disentangle or at least to bring out the mechanisms underlying the variability of the responses to drought for this widely distributed mite.

Under climate change, it is expected that mites will experience harsher drought episodes with environmental conditions leading to the selection of drought-adapted mites. In agriculture, intensification of damage in normally wet areas during the first years of drought (see Legrand et al., 2000 for example) will probably be limited by physiological costs and progressively lead to adaptation as suggested by the mite responses in the driest areas of this study. These uncertainties are important to consider for future strategies of pest management.

## Data, script and code availability

Data and statistical analysis are available here https://doi.org/10.15454/ICM8EF

## Acknowledgements

We thank Judit Linka (NIAB EMR, East Malling, United Kingdom), Lucía Adriana Escudero-Colomar (IRTA, Estació Experimental Agricola Mas Badia, Spain), Vassilis Litskas and Menelaos Stavrinides (Cyprus University of Technology, Cyprus) and Jean-Baptiste Thibord and Cyril Hannon (Arvalis - Institut du Végétal, France) who helped us for providing mites populations and Nathalie Smit (INRAE Montpellier) for her statistical advice.

We also thank Inês Fragata and Raul Costa-Pereira, the two PCI editors, Bastien Castagneyrol and two anonymous reviewers for their work and comments which greatly improved the manuscript.

## References

Auger P., Migeon A., Ueckermann E.A., Tiedt L., Navajas M. 2013. Evidence for synonymy between Tetranychus urticae and Tetranychus cinnabarinus (Acari, Prostigmata, Tetranychidae): Review and new data. Acarologia, 53: 383–415.

Bean D.W., Dalin P., Dudley T.L. 2012. Evolution of critical day length for diapause induction enables range expansion of Diorhabda carinulata, a biological control agent against tamarisk (Tamarix spp.). Evolutionary Applications, 5: 511–523.

Bendena W.G., Boudreau J.R., Papanicolaou T., Maltby M., Tobe S.S., Chin-Sang I.D. 2008. A Caenorhabditis elegans allatostatin/galanin-like receptor NPR-9 inhibits local search behavior in response to feeding cues. Proceedings of the National Academy of Sciences, 105: 1339–1342.

Chandler L.D., Archer T.L., Ward C.R., Lyle W.M. 1979. Influences of irrigation practices on spider mite 1. Densities on field corn 2. Environmental Entomology, 8: 196-201.

Chaves M.M., Maroco J.P., Pereira J.S. 2003. Understanding plant responses to drought - from genes to the whole plant. Functional Plant Biology, 30: 239–264.

Chen L., Sun J.-T., Jin P.-Y., Hoffmann A.A., Bing X.-L., Zhao D.-S., Xue X.-F., Hong X.-Y. 2020. Population genomic data in spider mites point to a role for local adaptation in shaping range shifts. Evolutionary Applications, 00: 1–15.

Crooker A. 1985. Embryonic and juvenile development. In: Helle W., Sabelis M.W., (Eds). Spider mites. Their biology, natural enemies and control. Elsevier. Amsterdam. p. 149–163.

Crozier R.H. 1985. Adaptative consequences of male-haploidy. In: Helle W., Sabelis M.W., (Eds). Spider mites. Their biology, natural enemies and control. Elsevier. Amsterdam. p. 201–222.

Deutsch C.A., Tewksbury J.J., Tigchelaar M., Battisti D.S., Merrill S.C., Huey R.B., Naylor R.L. 2018. Increase in crop losses to insect pests in a warming climate. Science, 361: 916–919.

English-Loeb G.M. 1989. Nonlinear responses of spider mites to drought-stressed host plants. Ecological Entomology, 14: 45–55.

Fellous S., Angot G., Orsucci M., Migeon A., Auger P., Olivieri I., Navajas M. 2014. Combining experimental evolution and field population assays to study the evolution of host range breadth. Journal of Evolutionary Biology, 27: 911–919.

Fick S.E., Hijmans R.J. 2017. WorldClim 2: new 1-km spatial resolution climate surfaces for global land areas. International Journal of Climatology, 37: 4302–4315.

Gillman J.H., Rieger M.W., Dirr M.A., Braman S.K. 1999. Drought stress increases densities but not populations of Two-spotted Spider Mite on Buddleia davidii ‘Pink Delight’. 34: 280.

Hamann E., Blevins C., Franks S.J., Jameel M.I., Anderson J.T. 2021. Climate change alters plant– herbivore interactions. New Phytologist, 229: 1894–1910.

Helle W., Bolland H.R. 1967. Karyotypes and sex-determination in spider mites (Tetranychidae). Genetica, 38: 43–53.

Hoffmann A.A. 2017. Rapid adaptation of invertebrate pests to climatic stress? Current Opinion in Insect Science, 21: 7–13.

Hummel I., Pantin F., Sulpice R., Piques M., Rolland G., Dauzat M., Christophe A., Pervent M., Bouteillé M., Stitt M. et al. 2010. Arabidopsis plants acclimate to water deficit at low cost through changes of carbon usage: An integrated perspective using growth, metabolite, enzyme, and gene expression analysis. Plant Physiology, 154: 357–372.

IPCC. 2021. Summary for Policymakers. In: Masson-Delmotte V., Zhai P., Pirani A., Connors S.L., Péan C., Berger S., Caud N., Chen Y., Goldfarb L., Gomis M.I. et al., (Eds). Climate change 2021: The physical science basis. Contribution of Working Group I to the Sixth Assessment Report of the Intergovernmental Panel on Climate Change. Cambridge University Press.

Krainacker D.A., Carey J.R. 1990. Effect of age at first mating on primary sex-ratio of the two-spotted spider mite. Experimental & Applied Acarology, 9: 169–175.

Legrand G., Wauters A., Muchembled C., Richard-Molard M. 2000. The common yellow spider mite (Tetranychus urticae Koch) (Acari: Tetranychidea) in sugarbeet in Europe: a new problem. 63e Congrès Institut International de Recherches Betteravières, Interlaken, Switzerland.

Lehmann P., Ammunét T., Barton M., Battisti A., Eigenbrode S.D., Jepsen J.U., Kalinkat G., Neuvonen S., Niemelä P., Terblanche J.S., Økland B, Björkman C. 2020. Complex responses of global insect pests to climate warming. Frontiers in Ecology and the Environment, 18: 141–150.

Litskas V.D., Migeon A., Navajas M., Tixier M.-S., Stavrinides M.C. 2019. Impacts of climate change on tomato, a notorious pest and its natural enemy: small scale agriculture at higher risk. Environmental Research Letters, 14: 084041.

Logan J.A., Wollkind D.J., Hoyt S.C., Tanigoshi L.K. 1976. An analytic model for description of temperature dependent rate phenomena in arthropods. Environmental Entomology, 5: 1133–1140.

Migeon A., Dorkeld F. 2019. Spider Mites Web: a comprehensive database for the Tetranychidae [Internet]. INRA; [cited]. Available from: http://www.montpellier.inra.fr/CBGP/spmweb

Migeon A., Tixier M.-S., Navajas M., Litskas V.D., Stavrinides M.C. 2019. A predator-prey system: Phytoseiulus persimilis (Acari: Phytoseiidae) and Tetranychus urticae (Acari: Tetranychidae): worldwide occurrence datasets. Acarologia, 59: 301-307.

Nikolova I., Georgieva N., Naydenova J. 2014. Development and reproduction of spider mites Tetranychus turkestani (Acari: Tetranychidae) under water deficit condition in soybeans Pesticidi i fitomedicina, 29: 187-195.

Olazcuaga L., Foucaud J., Deschamps C., Loiseau A., Claret J.-L., Vedovato R., Guilhot R., Sevely C., Gautier M., Hufbauer R. A, Rode N. O., Estoup A. 2022. Rapid and transient evolution of local adaptation to seasonal host fruits in an invasive pest fly. BioRxiv 03.01.482503; https://doi.org/10.1101/2022.03.01.482503

Oloumi-Sadeghi H., Helm C.G., Kogan M., Schoeneweiss D.F. 1988. Effect of water stress on abundance of two spotted spider mite on soybeans under greenhouse conditions. Entomologia Experimentalis et Applicata, 48: 85–90.

R Core Team R.C. 2018. R: A Language and Environment for Statistical Computing. Vienna, Austria: R Foundation for Statistical Computing.

Sabelis M.W. 1981. Biological control of two-spotted spider mites using phytoseiid predators. I. Modelling the predator-prey interaction at the individual level. Wageningen: Centre for Agricultural Publishing and Documentation.

Sadras V.O., Wilson L.J., Lally D.A. 1998. Water deficit enhanced cotton resistance to spider mite herbivory. Annals of Botany, 81: 273–286.

Santamaria M.E., Auger P., Martinez M., Migeon A., Castanera P., Diaz I., Navajas M., Ortego F. 2018. Host plant use by two distinct lineages of the tomato red spider mite, Tetranychus evansi, differing in their distribution range. Journal of Pest Science, 91: 169–179.

Santamaria M.E., Diaz I., Martinez M. 2018. Dehydration stress contributes to the enhancement of plant defense response and mite performance on barley. Front. Plant Sci. 9:458

Seagraves M.P., Riedell W.E., Lundgren J.G. 2011. Oviposition preference for water-stressed plants in Orius insidiosus (Hemiptera: Anthocoridae). J Insect Behavior, 24: 132–143.

Sengupta S., Cai W. 2019. A Quarter of Humanity Faces Looming Water Crises. The New York Times. New-York.

Showler A.T. 2013. Water deficit stress - Host plant nutrient accumulations and associations with phytophagous arthropods. In: Vahdati K., Leslie C., (Eds). Abiotic Stress - Plant Responses and Applications in Agriculture. London: IntechOpen. p. 387–410.

Sousa V.C., Zélé F., Rodrigues L.R., Godinho D.P., Charlery de la Masselière M., Magalhães S. 2019. Rapid host-plant adaptation in the herbivorous spider mite Tetranychus urticae occurs at low cost. Current Opinion in Insect Science, 36: 82–89.

Tomczyk A., Kropczyńska D. 1985. Effects on the host plant. In: Helle W., Sabelis M.W., (Eds). Spider mites. Their biology, natural enemies and control. Elsevier. Amsterdam. p. 317–329.

Trabucco A., Zomer R. 2019. Global Aridity Index and Potential Evapotranspiration (ET0) Climate Database v2 [Internet]. [cited]. Available from: https://figshare.com/articles/dataset/Global_Aridity_Index_and_Potential_Evapotranspiration_ET0_Climate_Database_v2/7504448

Van Petegem K.H.P., Boeye J., Stoks R., Bonte D. 2016. Spatial selection and local adaptation jointly shape life history evolution during range expansion. Am. Nat., 188: 485–498.

Venables W.N., Ripley B.D. 2002. Modern applied statistics with S. New-York: Springer.

Verslues P.E., Agarwal M., Katiyar-Agarwal S., Zhu J., Zhu J.-K. 2006. Methods and concepts in quantifying resistance to drought, salt and freezing, abiotic stresses that affect plant water status. The Plant Journal, 45: 523-539.

Walter J., Hein R., Auge H., Beierkuhnlein C., Löffler S., Reifenrath K., Schädler M., Weber M., Jentsch A. 2012. How do extreme drought and plant community composition affect host plant metabolites and herbivore performance? Arthropod-Plant Interactions, 6: 15–25.

Wickham H. 2016. ggplot2: Elegant graphics for data analysis. New York: Springer-Verlag.

Ximénez-Embún M.G., Castañera P., Ortego F. 2017a. Drought stress in tomato increases the performance of adapted and non-adapted strains of Tetranychus urticae. Journal of Insect Physiology, 96: 73-81.

Ximénez-Embún M.G., Glas J.J., Ortego F., Alba J.M., Castañera P., Kant M.R. 2017b. Drought stress promotes the colonization success of a herbivorous mite that manipulates plant defenses. Experimental and Applied Acarology, 73: 297–315.

Ximénez-Embún M.G., Ortego F., Castañera P. 2016. Drought-Stressed Tomato Plants Trigger Bottom–up effects on the invasive Tetranychus evansi. PLoS One, 11: e0145275.

Youngman R.R., Barnes M.M. 1986. Interaction of spider mites (Acari: Tetranychidae) and water stress on gas-exchange rates and water potential of almond leaves. Environmental Entomology, 15: 594-600.

Youngman R.R., Sanderson J.P., Barnes M.M. 1988. Life history parameters of Tetranychus pacificus McGregor (Acari: Tetranychidae) on almonds under differential water stress. Environmental Entomology, 17: 488-495.

