## Supplementary material for "The response to drought-stressed host plants varies among herbivorous mite populations from a climate gradient"

### 1 Climate data

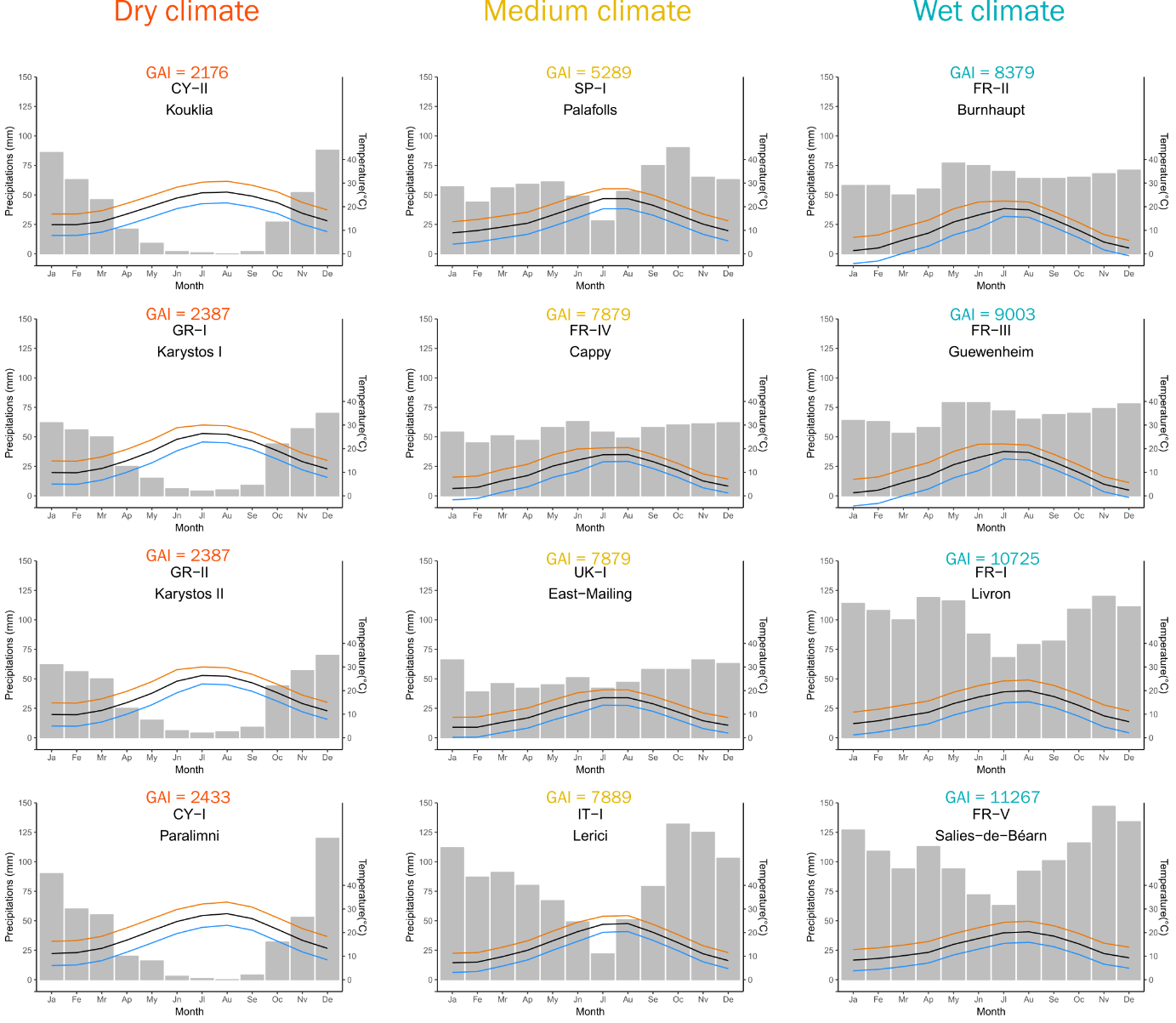

Figure S1. Gaussen ombro-thermic diagrams of the sampled locations. Grey bars: monthly precipitations (mm); blue lines: minimal monthly temperatures (°C); black lines: average monthly temperatures (°C); brown lines: maximal monthly temperatures (°C).

### 2 Watering control and drought stress assesment

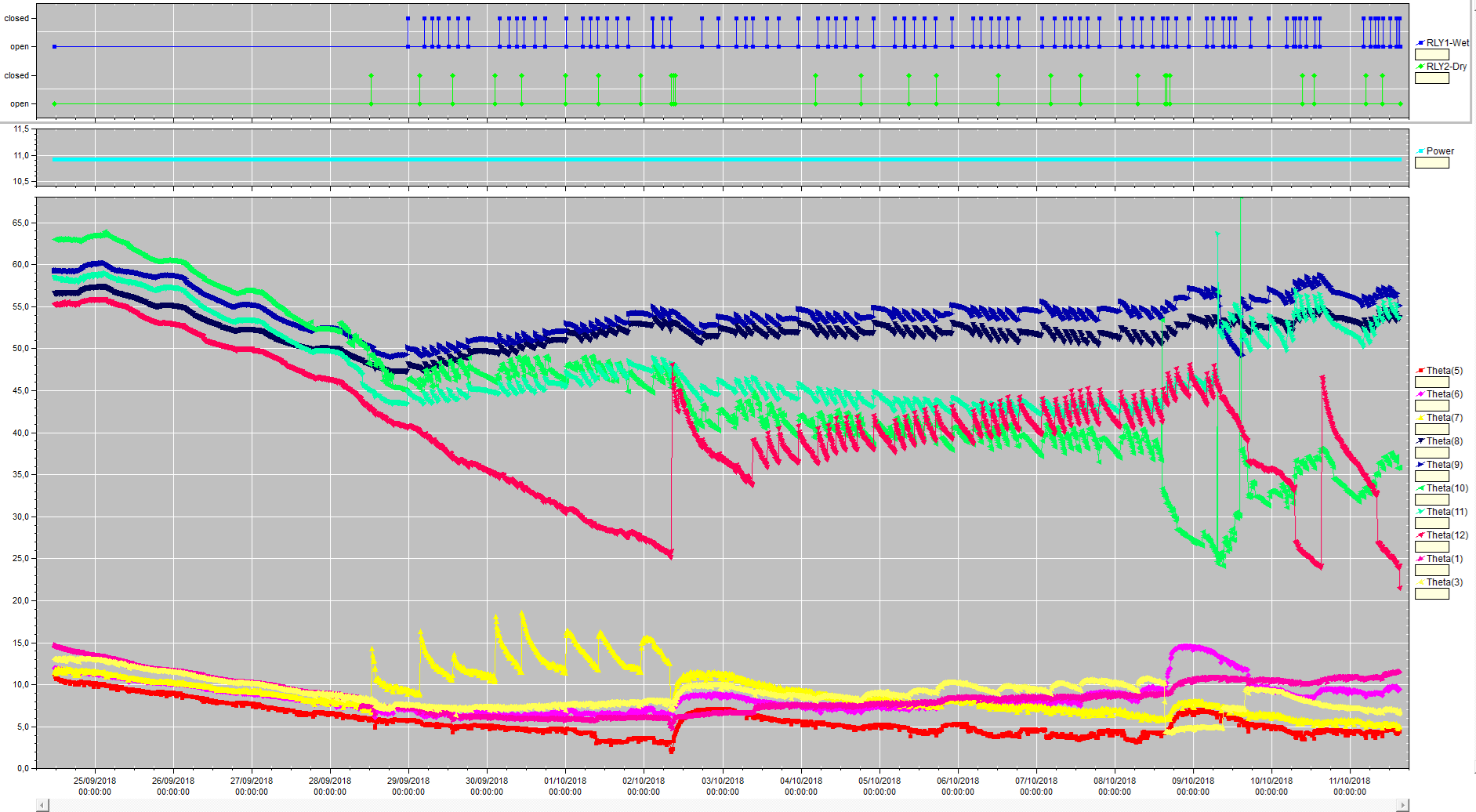

Well watered

Drought stressed

**Peat mix RH (%)**

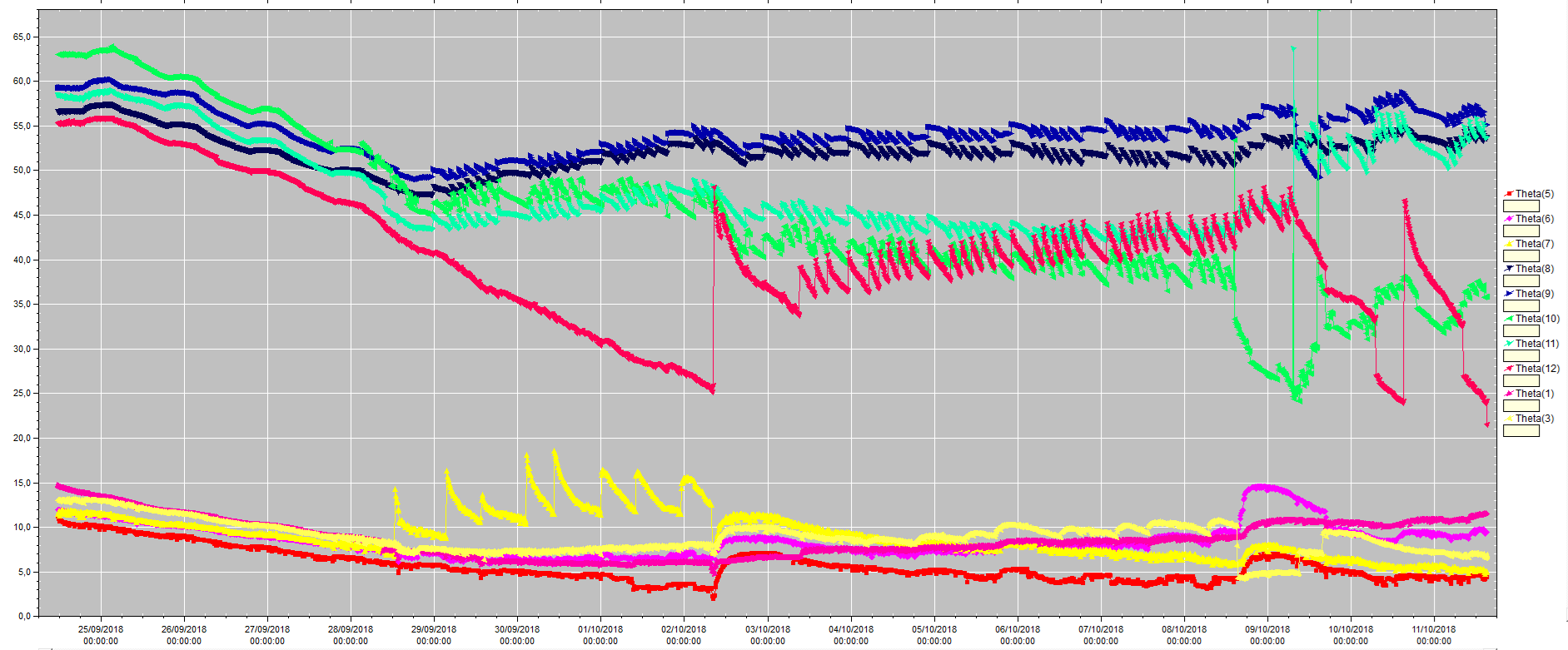

Well watered

Drought stressed

Watering events

Figure S2. Watering frequency (upper part) and soil moisture status (lower part) of bean plants of the experiment I in the well-watered and drought stressed modalities using automatic watering (5 moisture sensors SM150 per water treatment linked to GP2 Data Logger (Delta-T Devices Ltd) and to DeltaLINK 3.1.1 PC software).

Mite infestation

**Days after sowing (DAS)**

**Leaf stomatal conductance (mmol/m²s-1)**

|  | Well watered | Drought stressed |
| --- | --- | --- |
| 11 | 165.2 | 89.4 |
| 14 | 182.7 | 93.4 |
| 17 | 192.1 | 68.7 |
| 19 | 164.8 | 87.2 |
| 21 | 146.1 | 89.3 |
| 24 | 127.7 | 86.6 |
| 27 | 122.8 | 75.6 |

Figure S3. Water stress status (drought stressed and well-watered) of bean trial plants of the experiment I according to the leaf stomatal conductance. Leaf stomatal conductance was measured with seed leave from 10 to 19 days after sowing and with the central leaflet of the first trifoliate leaf from 21 days after sowing until the end of the experiment.

**Days after sowing (das)**

**Peat mix RH (%)**

**Pots weight (g)**

|  | Well watered | Drought stressed | Well watered RH | Drought stresses RH |
| --- | --- | --- | --- | --- |
| 10 | 1248.7 | 662.5 | 63.2 | 14.75 |
| 12 | 1300.3 | 598 | 70.4 | 11.75 |
| 14 | 1281.7 | 545.8 | 67.6 | 10.55 |
| 17 | 1156.7 | 469 | 55.4 | 9.5 |
| 19 | 1208.2 | 450.5 | 59.9 | 7.2 |
| 21 | 1173.6 | 424.7 | 60.5 | 11.65 |
| 24 | 998.4 | 393 | 49.1 | 6.55 |
| 26 | 1060.3 | 391 | 53.7 | 6.35 |
| 28 | 1036 | 374 | 54.3 | 6.75 |
| 31 | 837.8 | 342.2 | 35.6 | 7.37 |
| 33 | 933.1 | 343.4 | 46.2 | 6.3 |

Figure S4. Pots weight and peat mix RH of bean breeding plants of the experiment 2 in the well- watered and drought stressed modalities manually watered.

**Leaf stomatal conductance (mmol/m²s-1)**

**Days after sowing (das)**

|  | Well watered | Drought stressed |
| --- | --- | --- |
| 10 | 161.1 | 88 |
| 12 | 163.1 | 84.6 |
| 14 | 143 | 75.3 |
| 17 | 103.8 | 57.3 |
| 19 | 123 | 55.1 |
| 21 | 177.4 | 54 |
| 24 | 122.5 | 41.5 |
| 26 | 122.6 | 40.1 |
| 28 | 124.3 | 40.2 |
| 31 | 78.7 | 29.3 |
| 33 | 92 | 36.9 |

Figure S5. Water stress status (drought stressed and well-watered) of bean breeding plants in experiment 2 according to the leaf stomatal conductance. Leaf stomatal conductance was measured with seed leave from 10 to 33 days after sowing and with the central leaflet of the first trifoliate leaf from 21 days after sowing until the end of the experiment.

### 3 Confining area

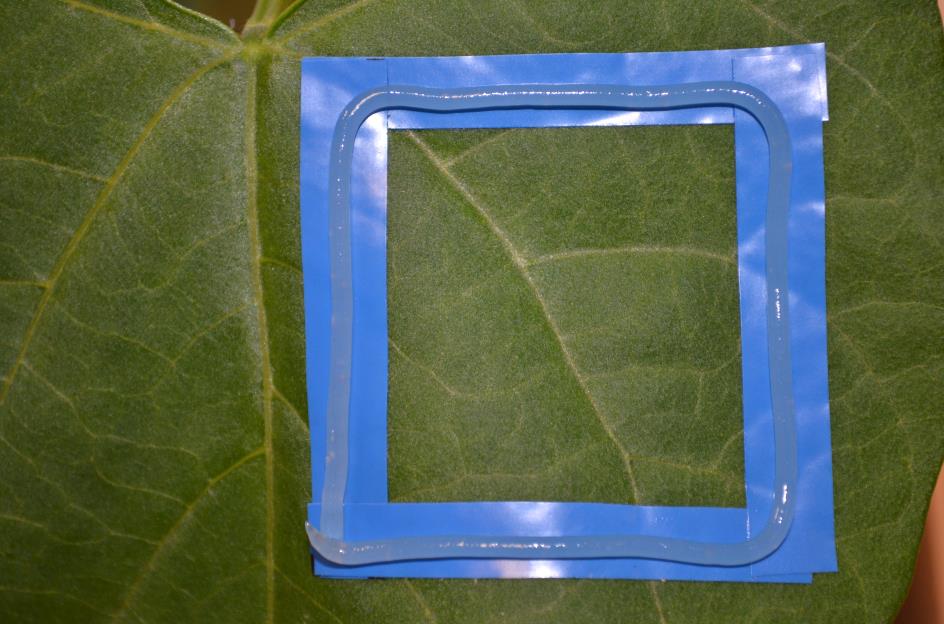

Figure S6. Mite confining arena built on bean cotyledon leaf. Each squared arena was delimited by four flexible PVC tape stripes (electrical insulation tape, Coteka, Chaulnes, France) of 50 x 7.5 mm forming an internal square about 18 cm² glued onto the leaf surface. To avoid mite losses, Vaseline (CAS 8009-03-8; Gifrer, Décines-Charpieu, France) mixed with 10% organic olive oil (Bio Classico, Carapelli Firenze SpA, Tavarnelle Val di Pesa, Firenze, Italy) was applied on the tape stripe to form a cord (that delimits a squared arena) that spider mites cannot cross.

### 4 Results

Table S1. Development time for each population of *Tetranychus urticae* on well-watered and drought stressed plants. Logit values of the 50% adult emergence +/- standard-error (in days). ANOVA analysis (*χ*^2^ model) to test the effect of watering regime on logistic regression: F-value, Df: degrees of freedom (10 points of comparison over time), *P*-value. Decrease of development duration on drought stressed plants is indicated by a downward arrow.

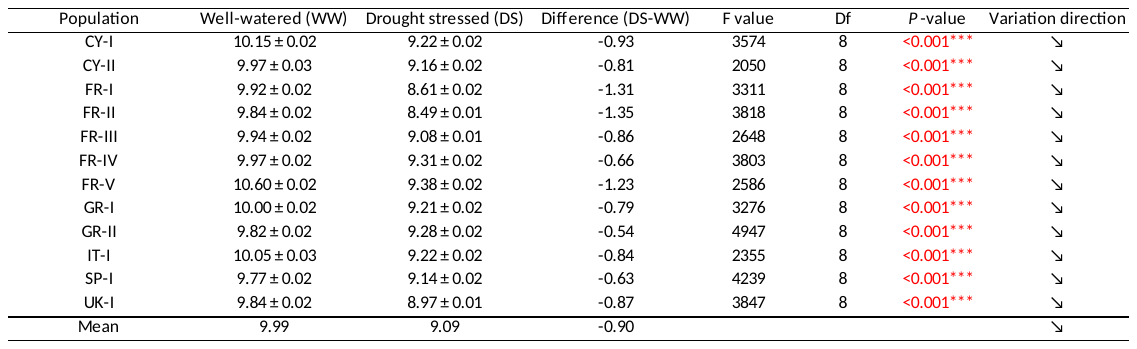

Table S2. Fecundity of the females for each population of *Tetranychus urticae* on well-watered and drought-stressed plants. Mean values +/- standard-erroor. *t*-test results (F-value, Df: degrees of freedom, *P*-value). Decrease of fecundity on drought stressed plants is indicated by a downward arrow.

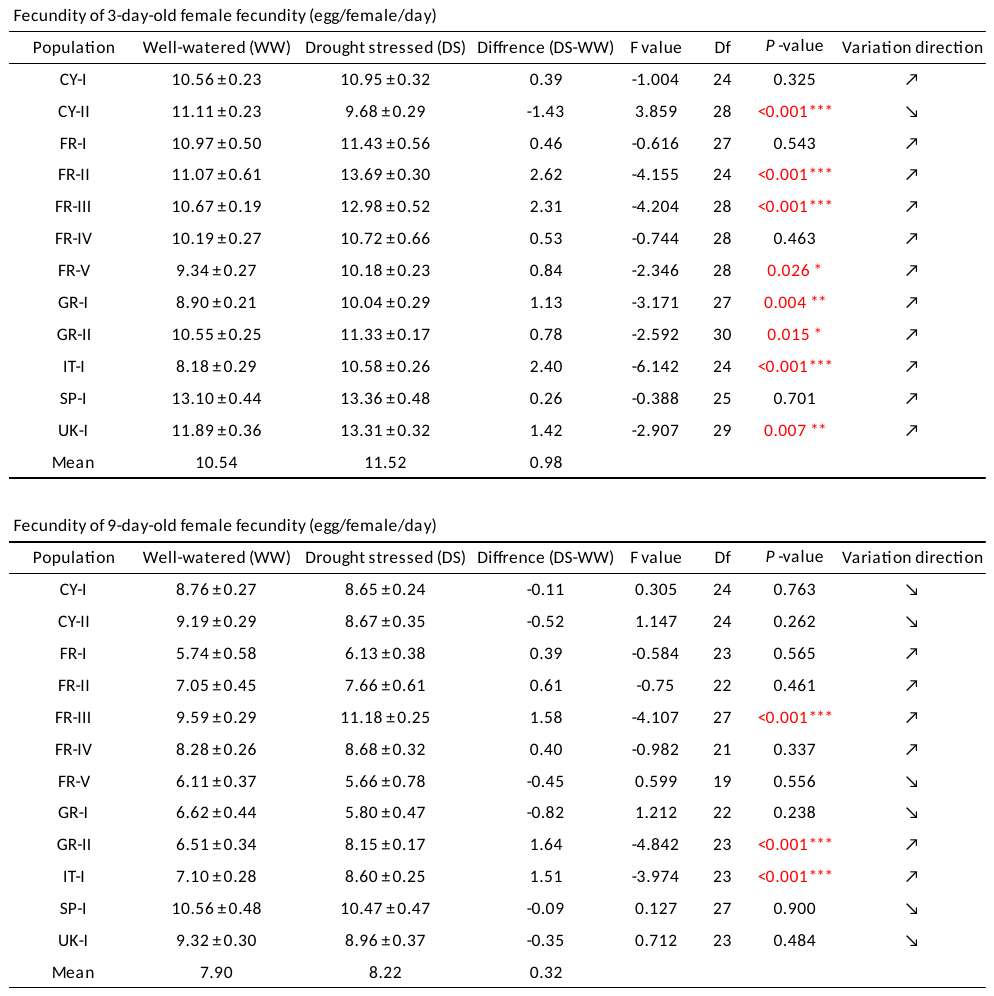

Table S3. Leaving rate of females for each population of *Tetranychus urticae* on well-watered and drought stressed plants. Mean values +/- standard-error. *t*-test results (F-value, Df: degrees of freedom, *P*-value). Decrease of leaving on drought stressed plants is indicated by a downward arrow.

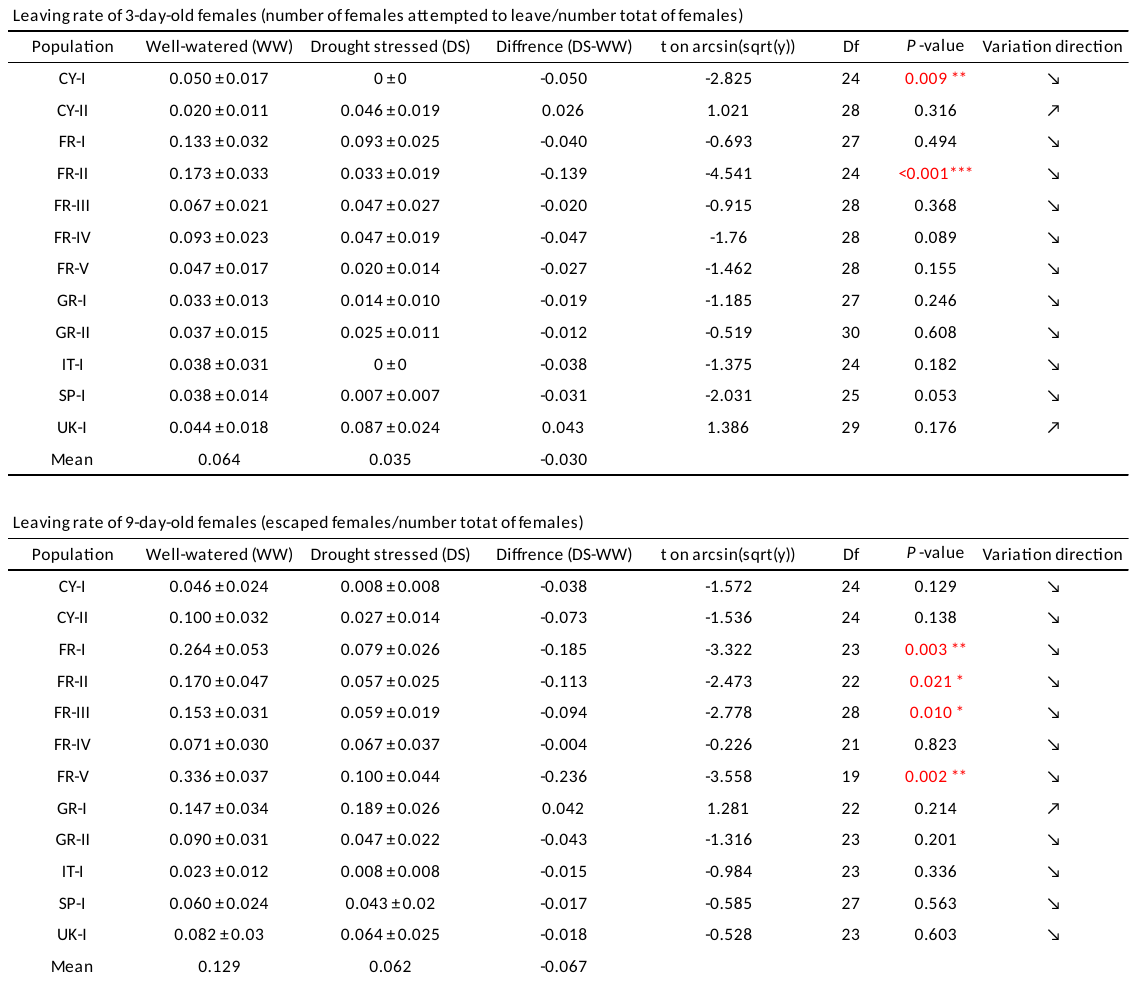

Table S4. Progeny sex-ratio for each population of *Tetranychus urticae* on well-watered and drought stressed plants (number of females / (number of males + females)). Mean values +/- standard-error. *t*-test results (F-value, Df: degrees of freedom, *P*-value). Decrease of sex-ratio on drought stressed plants is indicated by a downward arrow.

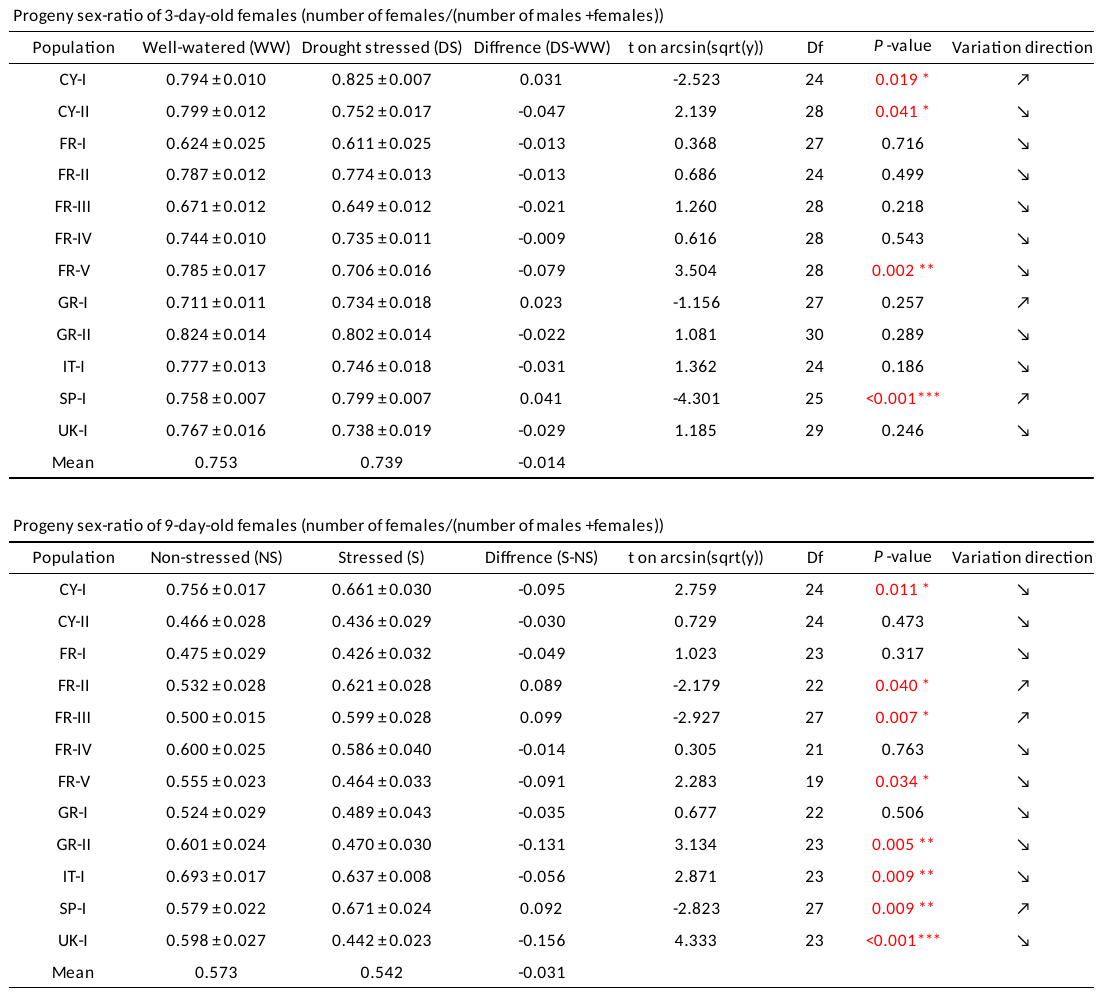
